# Revisiting the carbon isotope discrimination and water use efficiency relation: the influence of mesophyll conductance

**DOI:** 10.1101/2020.07.06.188920

**Authors:** Wei Ting Ma, Guillaume Tcherkez, Xu Ming Wang, Rudi Schäufele, Hans Schnyder, Yusheng Yang, Xiao Ying Gong

## Abstract

- The carbon isotope discrimination (Δ) has been used widely to infer intrinsic water-use efficiency (iWUE) of C_3_ plants, a key parameter linking carbon and water fluxes. Despite the essential role of mesophyll conductance (*g*_m_) in photosynthesis and Δ, its effect on Δ-based predictions of iWUE has generally been neglected.
- Here, we derive a mathematical expression of iWUE as a function of Δ that includes *g*_m_ (iWUE_mes_) and exploits the *g*_m_-stomatal conductance (*g*_sc_) relationship across drought-stress levels and plant functional groups (deciduous or semi-deciduous woody, evergreen woody and herbaceous species) in a global database. iWUE_mes_ was further validated with an independent dataset of online-Δ and CO_2_ and H_2_O gas exchange measurements with seven species.
- Drought stress reduced *g*_sc_ by 52% and *g*_m_ by 45% averaged over all plant functional groups, but had no significant effect on the *g*_sc_/*g*_m_ ratio, suggesting a well-constrained *g*_sc_/*g*_m_ ratio of 0.79±0.07 (95%CI, *n*=198) across plant functional groups and drought-stress treatments. Due in part to the synchronous behavior of *g*_sc_ and *g*_m_, *g*_m_ was negatively correlated to iWUE. Incorporating the *g*_sc_/*g*_m_ ratio in the iWUE_mes_ model significantly improved the estimation of iWUE compared to the simple model.
- The inclusion of *g*_m_ effects, even using a fixed *g*_sc_/*g*_m_ ratio of 0.79 when *g*_m_ is unknown, proved desirable to eliminate significant bias in estimating iWUE from Δ across various C_3_ vegetation types.

## INTRODUCTION

Water use efficiency (WUE) is a capital parameter of leaf physiology to assess photosynthetic performance and the trade-off between CO_2_ fixation and water loss. In addition to photosynthetic capacity, the key actor of WUE is stomatal conductance, which effectively plays a pivotal role in terrestrial carbon and water cycles because it constrains the diffusion of CO_2_ (*g*_sc_) and water vapor (*g*_sw_, with *g*_sw_=1.6*g*_sc_) between leaf photosynthetic tissues and the atmosphere. Leaf intrinsic water use efficiency (iWUE; see definitions in Table 1) is thus defined as the net photosynthetic rate (*A*_n_) per unit *g*_sw_ and is given by (disregarding boundary layer resistance):

**Table 1.** Definitions of abbreviations used for intrinsic water-use efficiency in this paper.

| Abbreviations | Definition |
| --- | --- |
| iWUE | Generic term for intrinsic water-use efficiency |
| iWUE<sub>obs</sub> | Observed value of intrinsic water use efficiency using gas exchange ($A_n/g_{sw}$) |
| iWUE<sub>sim</sub> | Predicted intrinsic water-use efficiency using the simplified model of $^{12}\text{C}/^{13}\text{C}$ discrimination (Eqn 2) |
| iWUE<sub>mes</sub> | Predicted intrinsic water-use efficiency using the model of $^{12}\text{C}/^{13}\text{C}$ discrimination accounting for mesophyll conductance but not boundary layer conductance, ternary effects nor day respiration (Eqn 12) |
| iWUE<sub>com</sub> | Predicted intrinsic water used efficiency using the model of $^{12}\text{C}/^{13}\text{C}$ discrimination accounting for all effects, including mesophyll conductance, ternary effects and day respiration (Eqn 10) |
| iWUE<sup>*</sup><sub>com</sub> | Predicted intrinsic water used efficiency using the model of $^{12}\text{C}/^{13}\text{C}$ discrimination accounting for all effects, including mesophyll conductance, ternary effects, boundary layer conductance and day respiration, and assuming that the definition of iWUE also includes boundary layer conductance in the denominator (Eqn 8) |

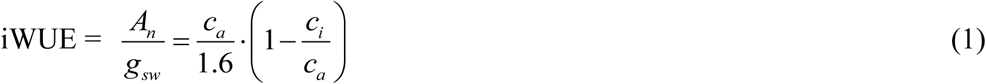

where *c*_a_ is the concentration of CO_2_ in the atmosphere and *c*_i_ the intercellular CO_2_ concentration. iWUE is useful to both describe plant physiological responses to environmental parameters (such as temperature and water availability) and understand the role of vegetation in hydrological cycles via its influence on water vapor and thus rainfall and runoff at regional to continental scales (Betts *et al*., 2007; Kooperman *et al*., 2018). As such, some researchers have called for more quantitative data on iWUE values in response to climate change to improve climate predictions (Adams *et al*., 2019, 2020). However, our ability to predict the response of carbon and water cycles in a changing climate depends ultimately on our understanding of mechanisms that control variation in iWUE.

Our current knowledge of iWUE is mostly based on its relationship with the photosynthetic ^12^C/^13^C isotope discrimination (Δ, Farquhar *et al*., 1982, 1989), which explicitly describes the isotopic fractionation associated with physical and biochemical processes during photosynthetic gas exchange. Using the simplified, linear relationship of Farquhar *et al*. (1982), neglecting mesophyll resistance (i.e. assuming an infinite internal conductance), the intrinsic water use efficiency (here denoted as iWUE_sim_, see Table 1 for definitions) is obtained from *c*_i_/*c*_a_ as:

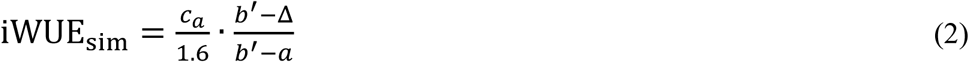

where *a* (4.4‰) is the ^12^C/^13^C discrimination during CO_2_ diffusion through stomata, *b’* (27‰) is the net discrimination during carboxylation (includes the potential contribution of effects other than Rubisco-catalysed reactions), and Δ is the net fractionation during photosynthesis that can be calculated from the isotope composition (δ^13^C) of atmospheric CO_2_ and plant fixed carbon (Farquhar & Richards, 1984; Ubierna *et al*., 2018). Δ can be estimated from on-line ^13^CO_2_/^12^CO_2_ gas exchange during photosynthesis or the δ^13^C of photosynthetic products (e.g. sugars and bulk leaf organic carbon). The iWUE_sim_ model (Eqn 2) has been widely used to infer iWUE from δ^13^C values in climate archives like herbage (Koehler *et al*., 2012), tree rings (Frank *et al*., 2013; van der Sleen *et al*., 2015), or animal tissues (Barbosa *et al*., 2010). Compared to instantaneous iWUE derived from gas exchange measurements, biomass-based Δ provides a time-integrated iWUE and enables time-series analysis spanning from days to millennia (Soh *et al*., 2019; Adams *et al*., 2020). This time-integrated iWUE is particularly useful for screening of high water-use efficiency crops thus has been used in breeding programs (for a review, see Flexas *et al*., 2013). Similarly, genetic determinants of iWUE have been searched for nearly 25 years using biomass-based Δ as a quantitative trait in QTL analyses (Rebetzke *et al*. 2008; Yang *et al*. 2016). Also, using surveys with many plant samples of diverse origins, spatial variation in vegetation iWUE could be assessed or integrated at a regional (Wittmer *et al*., 2010) or global scale (Cornwell *et al*., 2018; Adams *et al*., 2020). More recently, δ^13^C-derived iWUE, together with transpiration inferred from sap flow, have been used to estimate canopy photosynthesis (Klein *et al*., 2016). However, it has been argued that the iWUE_sim_ model gives biased predictions of iWUE, limiting its application to qualitative assessments (Seibt *et al*., 2008; Barbosa *et al*., 2010).

In fact, the main limitation of the use of iWUE_sim_ is the simplifying assumption that mesophyll conductance (*g*_m_, the diffusive conductance of CO_2_ from intercellular space to the site of carboxylation) is infinite (Seibt *et al*., 2008; Franks *et al*., 2013; Stangl *et al*., 2019). This assumption, theoretically, leads to an overestimation of iWUE (see Theory below) because *g*_m_ is not conservatively high (for reviews see Flexas *et al*., 2008, 2012) and thus this causes an increased CO_2_ drawdown from the atmosphere to chloroplasts and thereby decreases the actual isotope fractionation Δ. Furthermore, responses of *g*_m_ to environmental stimuli have been reported, including water stress (Galmes *et al*., 2007; Warren *et al*., 2011; Barbour & Kaiser, 2016), temperature (Yamori *et al*., 2006; Evans & Von Caemmerer, 2013), and CO_2_ mole fraction (Flexas *et al*., 2007; Tazoe *et al*., 2011). In other words, applying either an infinite or a species-specific, constant *g*_m_ in iWUE models is perhaps improper. Water stress is particularly relevant in the context of water utilization by plant leaves, because it strongly influences *g*_sw_ and thus iWUE (Lin *et al*., 2015). However, its impact on *g*_m_ is currently uncertain. Some studies have reported that both *g*_m_ and *g*_sw_ decreased under edaphic drought, i.e. low soil water content (Galmes *et al*., 2007; Cano *et al*., 2014). In addition, while stomatal limitation of photosynthetic gas exchange has been found to be the dominant factor in the early stage of drought, the decrease in *g*_m_ further restricts photosynthesis later on (Perez-Martin *et al*., 2014). Vapor pressure deficit (VPD), a key factor that influences both plant water potential and stomatal conductance, has been found to decrease (Loucos *et al*., 2017) or have no influence on *g*_m_ (Warren, 2008; Stangl *et al*., 2019). In principle, a low sensitivity of *g*_m_ to water deficit is beneficial for maintaining a high CO_2_ assimilation rate (Cano *et al*., 2014). That is, a combination of a low g_sw_ and a high *g*_m_ should lead to a high iWUE if photosynthetic capacity is maintained (Barbour *et al*., 2010; Flexas *et al*., 2013; Barbour & Kaiser, 2016). Accordingly, *g*_m_ has been suggested to be an important target for crop improvement (Cano *et al*., 2014; Barbour & Kaiser, 2016), although this hypothesis was not supported by experimental results of wheat genotypes (Barbour & Kaiser, 2016). Therefore, the responses of *g*_m_ and *g*_sw_ to long-term drought stress is not well understood, and the implication for iWUE estimation is not clear.

Having said that, incorporating mesophyll conductance is still challenging because measuring *g*_m_ involves technically demanding methods, such as online carbon isotope discrimination (Evans *et al*., 1986), chlorophyll fluorescence (Harley *et al*., 1992), or ^18^O mass balance (Gauthier *et al*. 2018). This explains why *g*_m_ measurements cannot be carried out at high-throughput and therefore, our knowledge of mechanisms driving variation in *g*_m_ is still limited. In practice, there is presently no available method to incorporate *g*_*m*_ effects in iWUE and carbon cycle models derived therefrom (Flexas *et al*., 2012; Sun *et al*., 2014a). Alternatively, empirical estimates can be used to scale *g*_m_ to other photosynthetic parameters, since *g*_m_ has been shown to be positively related to net assimilation rate or *g*_sc_ (Flexas *et al*., 2008; Gong *et al*., 2018). Such relationships could be useful – providing they are well-constrained – for improving estimates of iWUE. However, such relationships have not been tested with a global compiled dataset of a diverse range of functionally distinct species.

Taken as a whole, the broad application of ^12^C/^13^C isotope fractionation to compute intrinsic water use efficiency generally neglects the impact of mesophyll conductance, either because of technical reasons (difficult access to mesophyll conductance measurements) or because its impact is assumed to be relatively modest (assumption of infinite *g*_m_ in iWUE_sim_). However, the reliability of iWUE estimates omitting mesophyll conductance is questionable, in particular when the photosynthetic limitation associated with mesophyll resistance is large. Here, we derive an expression of iWUE that includes the effect of *g*_m_ (denoted as iWUE_mes_), and test the numerical impact of *g*_m_ using independent experimental datasets. We address the following questions: 1) can we substantially improve the prediction of iWUE (compared with iWUE_sim_) by including a term accounting for *g*_m_? and, 2) could we take advantage of a *g*_m_-*g*_sc_ relationship and if so, can it be generalized to compute iWUE? To answer these questions, we collated a database of more than 300 pairs of *g*_sc_ and *g*_m_ measurements of more than 84 species from >40 publications. We analyzed the response of *g*_m_ and *g*_sc_ to long-term drought stress in different plant functional groups to parameterize iWUE_mes_. To test whether iWUE_mes_ provides reliable estimates of intrinsic water use efficiency, we used an independent dataset of iWUE_obs_ values (obtained directly from gas exchange), and on-line photosynthetic Δ corrected for side effects of respiratory isotopic imbalance (Gong *et al*., 2015). The latter dataset compiled measurements obtained with seven species grown in controlled-environment conditions.

## MATERIALS AND METHODS

### Theory

Farquhar *et al*. (1982) and Farquhar & Cernusak (2012) established a comprehensive theoretical model to describe the photosynthetic ^13^C discrimination of C_3_ leaves (Δ) as:

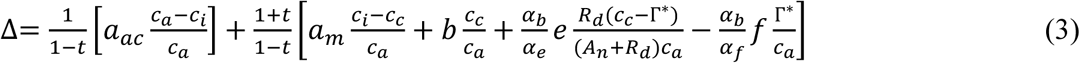

where *c*_a_, *c*_s_, *c*_i_ and *c*_c_ are CO_2_ mole fractions in the atmosphere, at the leaf surface, and in the substomatal cavity and chloroplast, respectively. *a*_m_ (1.8‰) is the fractionation associated with CO_2_ dissolution and diffusion in the mesophyll; *R*_d_ represents the day respiration rate; Г* is the CO_2_ compensation point in the absence of mitochondrial respiration.; *b* (29‰), *e* (−6-0‰) and *f* (11‰) represent the fractionations due to carboxylation, daytime mitochondrial respiration and photorespiration, respectively. Also, *α*_*b*_=1+*b, α*_*e*_=1+*e, α*_*f*_=1+*f. t* is the ternary correction factor and *a*_ac_ is the weighted fractionation for diffusion across the boundary layer and stomata. They can be calculated as:

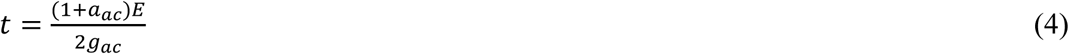

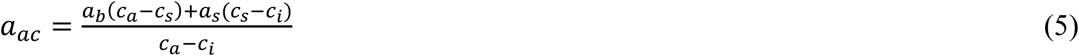

where *a*_b_ (2.9‰) and *a*_s_ (4.4‰) are the ^12^C/^13^C fractionations during CO_2_ diffusion through the leaf boundary layer and stomata, respectively. *E* is the transpiration rate. *c*_s_ is the CO_2_ concentration at the leaf surface; *g*_ac_ is the combined boundary layer and stomatal conductance to CO_2_.

From this point, we can use Δ to calculate an exact value of intrinsic water use efficiency that accounts for both internal conductance and ternary effects. It will be denoted as iWUE_com_. We recall here that accounting for ternary effects is so that:

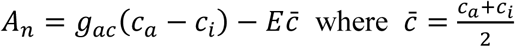

At this stage, there are two possibilities to compute intrinsic water use efficiency: accounting for boundary layer conductance, or disregarding boundary layer resistance. As per Eqn 3 to 5, the observed isotope fractionation accounts for boundary layer effects and the CO_2_ drawdown from *c*_*a*_ to *c*_*i*_ is the result of both boundary layer and stomatal resistance.

- If the definition of intrisic water use efficiency encapsulates both boundary layer and stomatal effects (we denote this iWUE as iWUE_com_^*^), we have:

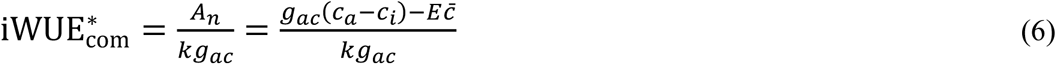

Where *k* is the coefficient between CO_2_ and H_2_O conductances (it equals 1.6 in Eqn 1). Using the expression *E* = *kg*_*ac*_*W* (where *W* is the water vapor drawdown from intercellular spaces to air), we have:

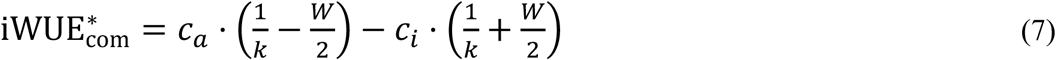

Substituting Eqns 6 and 7 into 3, we have (for intermediate calculations, see Note S1):

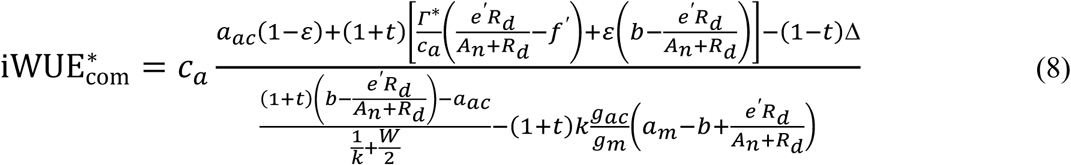

where *e’* = *eα*_*b*_*/α*_*e*_, *f’* = *fα*_*b*_*/α*_*f*_ and

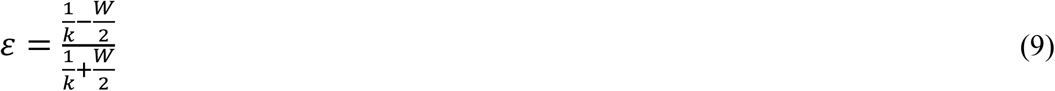

Here, it is useful to indicate the order of magnitude of terms found in Eqn 8 but not found in simplified expressions commonly used (such as Eqn 2). Assuming that *E* is typically about 2 mmol m^-2^ s^-1^ and stomatal conductance for CO_2_ is 0.1 mol m^-2^ s^-1^ gives *t* in the order of 1/100^th^ thus its impact is small. *W* is homogenous to a mole fraction and thus about 0.01 (i.e. typically in the order of 10 mmol H_2_O mol^-1^) and thus *ε* ≈ 0.98 and its impact is rather small. Taken as a whole, the ternary correction affects iWUE by about 1%.

- If intrinsic water use efficiency does not encapsulate boundary layer effects (it is then denoted as iWUE_com_), then:

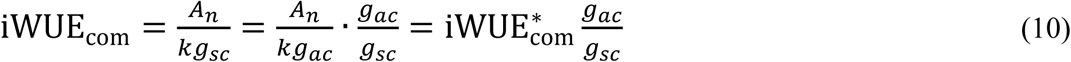

Note that Eqn 10 means that boundary layer effects are so that iWUE_com_ < iWUE^*^ _com_ since *g*_*ac*_ < *g*_*sc*_. When ternary corrections are ignored and boundary layer conductance is presumed to be infinite, then *t*=0, *ε*=1, *c*_a_=*c*_s_ and thus *a*_ac_=*a*_s_ and *g*_ac=_*g*_sc._ Eqn 8 simplifies to:

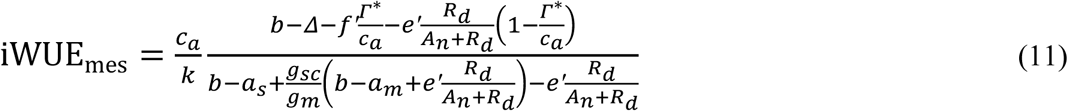

Here, the subscript “mes” indicates that this expression accounts for mesophyll conductance effects but not ternary correction or boundary layer effects. If the impact of day respiration is assumed to be negligible (*e* = 0), or if the photosynthetic fractionation has been corrected for *R*_d_ (see below), Eqn 11 gives the proxy:

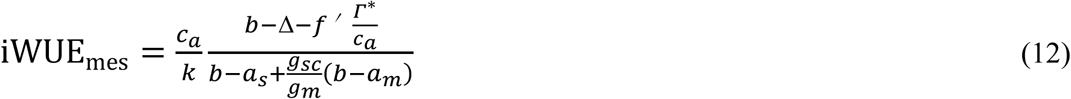

The simplifying assumptions used here to obtain Eqn 12, which have been commonly applied (Ubierna *et al*., 2018), are supported by sensitivity tests (see below). Compared with iWUE_sim_ (Eqn 2), iWUE_mes_ (Eqn 12) takes into account the effects of mesophyll conductance and photorespiration. Combining Eqn 12 and Eqn 2, iWUE_mes_ is related to iWUE_sim_ as:

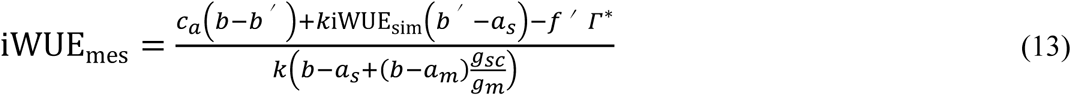

Eqn 12 can also be used to examine the sensitivity of iWUE_mes_ with respect to the *g*_sc_-to-*g*_m_ ratio. Using the partial derivative of iWUE_mes_ gives the elasticity (relative change in iWUE_mes_ for a unit variation in *g*_sc_/*g*_m_):

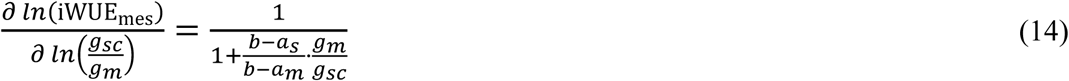

To assess the sensitivity of iWUE models (Eqn 8, 10 and 12) to different simplifications, we used theoretical data and varied the photosynthetic discrimination within 18 and 22‰ at different *g*_sc_/*g*_m_ ratios: *g*_sc_/*g*_m_=0 (iWUE_sim_ model), *g*_sc_/*g*_m_=0.4, 0.8 and 1.6 (iWUE_com_ and iWUE_mes_ models) (for details on calculations, see Table S1).

### Collection of literature data of g_m_ and g_sc_

To parameterize *g*_sc_/*g*_m_ in Eqn 12, we performed an extensive literature review of leaf-level *g*_m_ and *g*_sc_ measurements by searching the keywords ‘mesophyll conductance’ and ‘internal conductance’ in ISI Web of Science and checking the cited references in recent review articles of this topic. Data published before 2020 that met the following criteria were included in our analyses: (i) leaf-level *g*_m_ and *g*_sc_ data were reported; (ii) *g*_m_ was measured with either the online carbon isotope discrimination (Evans *et al*. 1986) or fluorescence (Harley *et al*., 1992) methods; and (iii) both control and edaphic drought treatments were included. We neglected results with short-term drought treatment (i.e. exposure time shorter than 2 days) and results of re-watering experiments, since acclimated parameters are more representative for the steady-state physiological status of plants compared with dynamically changing parameters. Gas exchange parameters (e.g. net assimilation rate, transpiration rate) were also collected if presented. Furthermore, we eliminated outliers which were defined by the criteria: *A*_n_/*g*_sc_>300, *g*_sc_/*g*_m_<0.2, or *g*_m_>0.8 mol m^-2^ s^-1^ (cf. Flexas *et al*., 2013). Average values for of each species in each treatment combination were compiled, and the compiled dataset included 198 pairs of *g*_m_ and *g*_sc_ measurements of 32 species under manipulated soil water availability or VPD from 15 studies (‘drought-stress dataset’). Additionally, published data reporting long-term treatments other than edaphic drought (e.g. temperature, salinity, soil nutrients, light, etc.) were collected and screened with the same criteria to detect outliers. We thus obtained a more comprehensive dataset (‘the main dataset’) that included 364 pairs of *g*_m_ and *g*_sc_ measurements of 84 species from 50 studies. We used the ‘drought-stress dataset’ to analyze the effects of water stress treatments and plant functional groups, and ‘the main dataset’ was used to examine the *g*_sc_-*g*_m_ relation in a larger dataset.

Statistical analyses on the compiled drought-stress dataset of *g*_m_, *g*_sc_, *A*_n_, and *g*_sc_/*g*_m_ were performed by using the general linear model of SPSS Statistics 19 (IBM Corp). The general linear model included water stress and plant functional groups (PFGs) and their interaction as the main factors. The factor of drought stress had two levels: control (non-stressed plants) and stressed. Plant species were classified into three PFGs: deciduous or semi-deciduous woody (DW), evergreen woody (EW), and herbaceous (HB, including annual, bi-annual and perennial) species. The parameters were logarithmic and square-root transformed to improve the normality before ANOVA tests or linear regressions, and both methods had no effect on the statistical results. We present here square-root transformed results because it improved the normality to a greater extent compared to the logarithmic transformation, as detected from histograms.

### Experimental data and photosynthetic ^13^C discrimination

To validate the iWUE_mes_ model (Eqn 12), we used independent datasets from two experiments (Gong *et al*. 2015, 2018) which included simultaneous measurements of photosynthetic gas exchange and online ^13^Cdiscrimination on fully expanded young leaves of seven species (Table 2). Most of the data have been published with the exception of *Vicia faba* and *Glycine max* grown and measured in the same way as the other species (Gong *et al*., 2018). These datasets were particularly suitable for evaluating the relation between Δ and iWUE because advanced protocols were applied to minimize any effect of diffusive leaks in the leaf cuvette and the isotopic disequilibrium between photosynthesis and day respiration. That system used a ^13^CO_2_/^12^CO_2_ gas exchange and labelling system connected with a portable leaf gas exchange apparatus (LI-6400, LICOR Inc., Lincoln, USA) and gas exchange mesocosms, with the air supply to the LI-6400 and mesocosms provided by mixing CO_2_-free, dry air and CO_2_ of known δ^13^C (Schnyder *et al*., 2003). This setup enabled the quantification of diffusive leak coefficients during measurements with intact leaves by measuring and manipulating ^13^CO_2_ and ^12^CO_2_ gradients between leaf cuvette and the surrounding air in the dark, thus effectively correcting leak artefacts (Gong *et al*. 2015, 2018).

**Table 2.** Overview of the two experimental datasets of Δ_obs_ in this study.

| Source | Gong *et al.* (2015) | Gong *et al.* (2018) |
| --- | --- | --- |
| Plant species | *Holcus lanatus* | *Hordeum vulgare*, *Triticum aestivum*, *Ricinus communis*, *Phaseolus vulgaris*, *Glycine max*, *Vicia faba* |
| Leaf types | Fully expanded young leaves | Fully expanded young leaves |
| Growth condition | Growth chamber, non-stressed condition | Growth chamber, non-stressed condition |
| Measurement condition | PPFD: 1600 $\mu\text{mol m}^{-2} \text{s}^{-1}$;<br>Leaf temperature: 25 °C;<br>[CO<sub>2</sub>]: 390 ppm; rH: 70% | PPFD: 700 $\mu\text{mol m}^{-2} \text{s}^{-1}$;<br>Leaf temperature, 23 °C<br>[CO<sub>2</sub>]: 390 ppm; rH: 70% |
| iWUE | Directly measured | Directly measured |
| Diffusive leak | Quantified and corrected | Quantified and corrected |
| Contribution of day respiration to $\Delta_{\text{obs}}$ | Quantified and corrected | Quantified and corrected |

The iWUE_mes_ model (Eqn 12) excludes the fractionation component associated with *R*_d_, therefore, to precisely compare the modeled and the observed iWUE, the isotopic fractionation of *R*_d_ (Δ_Rd_) should be subtracted from the observed net discrimination (Δ_AN_) as Δ_P_ =Δ_AN_ + Δ_Rd_, where Δ_P_ is the photosynthetic discrimination corrected for day respiration. Alternatively, our experimental data can be used to parameterize Eqn 11, and in that case Δ_Rd_ is not subtracted. Subtracting the effect of *R*_d_ is particularly useful when there is an isotopic disequilibrium whereby respiratory substrates have an isotope composition that is substantially different from that of current assimilates.

If we neglect ternary corrections in Eqn 3, the *R*_d_ corrected photosynthetic discrimination (Δ_P_) can be calculated as:

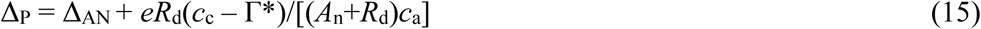

This equation is difficult to apply in practice because (i) the term *R*_d_ in Eqn 3 is the gross respiration rate (before refixation by photosynthesis and anapleurotic reactions) and is not easily measurable (Tcherkez *et al*. 2017); (ii) it requires the knowledge of *c*_c_ thus we would need to measure *g*_m_ first. We thus took advantage of the isotopic disequilibrium approach of Gong *et al*. (2015; 2018) to subtract the term associated with *R*_d_ from Δ_AN_. Assumptions on the isotope effect of *R*_d_ are further detailed in Note S2. Online discrimination measurements (Evans *et al*., 1986) were performed using a ^13^C-enriched CO_2_ source, followed by the measurement using a ^13^C-depleted CO_2_ source, so that the measured net assimilation rate (*A*_n_) could be partitioned into net photosynthesis in the absence of day respiration (*P*) and day respiration (*R’*_d_) (Gong *et al*., 2015). Applying isotopic mass balance, the photosynthetic discrimination (Δ_P_) was calculated from the measured net discrimination (Δ_AN_) as (Tcherkez *et al*., 2011; Gong *et al*., 2018):

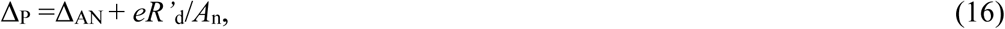

where *R’*_d_ is day respiration measured by the method of Gong *et al*. (2015; 2018) and *e* is the measured isotope fractionation of *R*_d_ relative to *A*_n_ determined as *e*=(δ_An_ – δ_Rd_)/(1+ δ_Rd_). It is assumed that δ_Rd_ was equal to the measured isotopic signature of leaf respiration in the dark (Gong *et al*. 2015, 2018). Importantly, Eqn 16 assumes that CO_2_ evolved by day respiration has a δ^13^C value that is distinct to that of photosynthesis. In other words, the calculation treats *R’*_d_ as a flux metabolically disconnected from recent photosynthesis during a short-duration of measurements (Note S2), and the related assumptions have been validated and discussed previously (Gong *et al*. 2015, 2017, 2018) and supported by other work (Barbour *et al*., 2017; Busch *et al*., 2020). Furthermore, taking different assumptions concerning the value of respiratory fractionation (Eqn 3 vs. Eqn S1) eventually had very minor influence on iWUE estimates (Note S2, Fig. S4).

Knowing Δ_P_, iWUE was calculated from the iWUE_mes_ model (Eqn 12) using a mean *g*_sc_/*g*_m_ of 0.79 derived from the compiled literature dataset (see below), and Г* as estimated from leaf temperature (*T*) as Г*=42.7+1.68(*T*-25)+0.012(*T*-25)^2^ following Brooks & Farquhar (1985). In addition, iWUE was also calculated from the simple model (iWUE_sim_ Eqn 2). The estimates of iWUE_mes_ and iWUE_sim_ were compared with the measured iWUE from gas exchange measurements (iWUE_obs_). The Root Mean Square Error (RMSE) was calculated to evaluate the prediction errors for the simple and the iWUE_mes_ model. The value of *g*_m_ of these plants was determined using the improved online Δ method and were published previously (Gong *et al*. 2015; 2018), except for the *Vicia faba* and *Glycine max* data. The detailed description of methods and equations for *g*_m_ are presented in Gong *et al*. (2015; 2018).

## RESULTS

### Sensitivity of iWUE models to g_m_ and simplifying assumptions

The expression of iWUE_mes_ (Eqn 12) includes the *g*_sc_/*g*_m_ ratio, implying in principle that iWUE_mes_ varies with *g*_m_. Accordingly, the expression of elasticity (Eqn 14) shows that estimating iWUE_mes_ from Δ using Eqn 12 is sensitive to the *g*_sc_/*g*_m_ ratio when *g*_m_ is varied in relation to *g*_sc_. For example, as is shown below, with a very high value of *g*_sc_/*g*_m_, e.g. *g*_sc_/*g*_m_ = 3, the elasticity value is 0.77. This implies a 77% change in iWUE_mes_ when *g*_sc_/*g*_m_ changes by 100%. By contrast, when *g*_m_ is large, the denominator in Eqn 14 is large and the elasticity value becomes small. Eqn 14 thus reflects the intuitive result that when *g*_m_ is large, its impact on Δ is small and thus numerically, the impact of the *g*<u>_sc_</u>/*g*_m_ ratio on iWUE is modest (and effectively nil, when *g*_m_ is infinite, as assumed by iWUE_sim_).

A sensitivity analysis was performed to quantify the effect of *g*_sc_/*g*_m_ ratio, dark respiratory fractionation *e*, photosynthetic fractionation *f* and ternary corrections on iWUE, as predicted by the different iWUE models (Fig. 1). Both iWUE_com_ and iWUE_mes_ were sensitive to the *g*_sc_/*g*_m_ ratio. The simplified model (that assumed *g*_sc_/*g*_m_=0) predicted the largest iWUE, as expected (Fig. 1a). At all *g*_sc_/*g*_m_ ratios, iWUE_com_ and iWUE_mes_ predictions were numerically very similar (with an error of generally less than 3 μmol mol^-1^, Fig. 1a). Furthermore, assuming an *f* of 0 or 20‰ led to a variation of predicted iWUE of about 5 μmol mol^-1^ (Fig. 1b). Similarly, ignoring respiratory fractionation or ternary correction caused an error of less than 2 μmol mol^-1^ (Fig. 1cd). Also, manipulating boundary layer conductance between 1 mol m^-2^ s^-1^ and infinite had very little influence on iWUE estimates (error less than 2 μmol mol^-1^, data not shown). Thus, the assumptions we used to derive iWUE_mes_ (Eqn 12) seem to be appropriate.

**Fig. 1.**
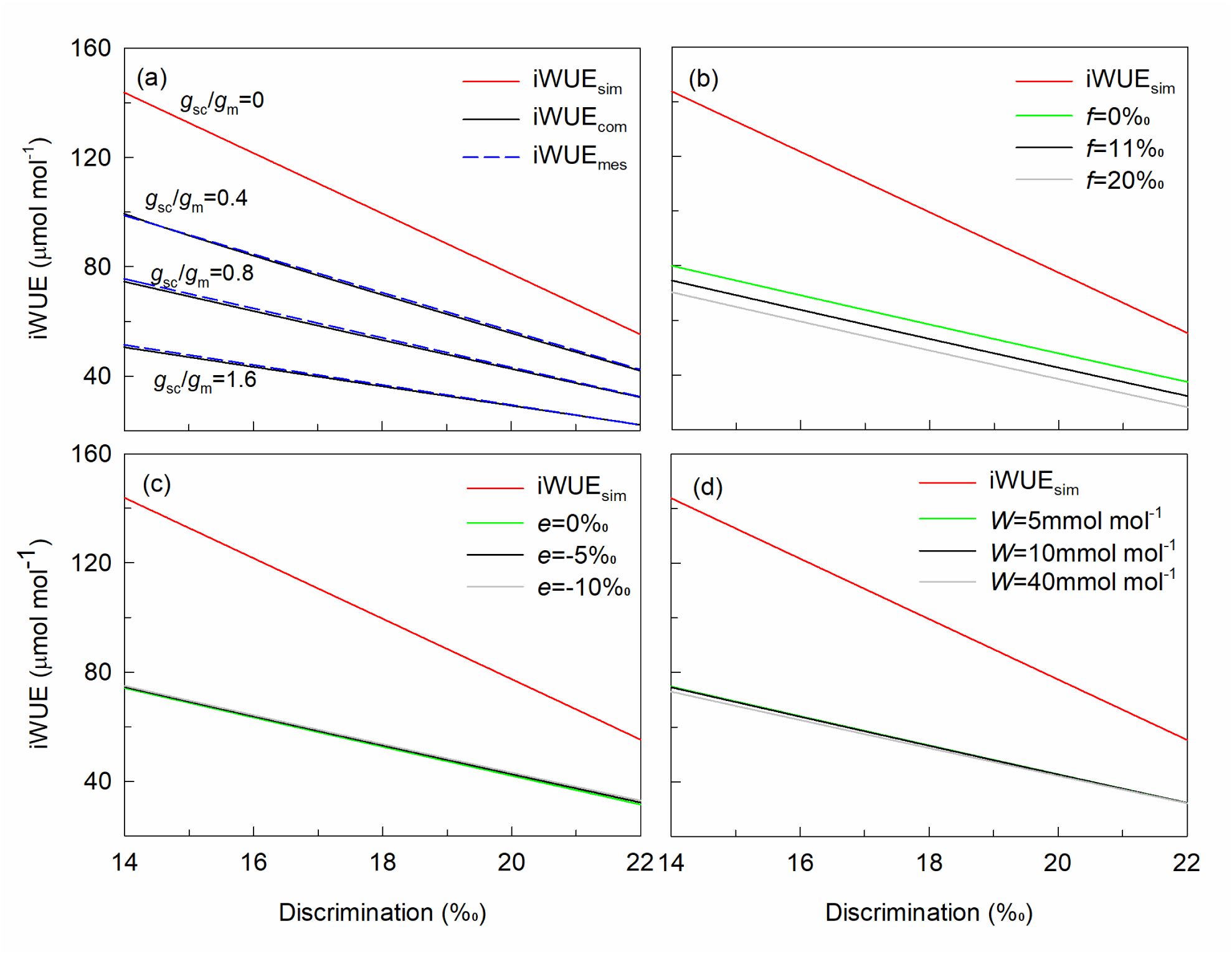
Sensitivity test of iWUE prediction by the iWUE_com_ model and the iWUE_mes_ model as affected by *g*_sc_/*g*_m_ ratios (a) and the influence of photorespiratory fractionation (b), mitochondrial respiratory fractionation (c) and leaf-to-air vapor concentration difference (*W*, mmol mol^-1^, d) on iWUE_com_ estimations. The black lines in the panels b-d represent the ‘standard scenario’ for data simulation with *g*_sc_/*g*_m_=0.8, *f*=11‰, *e*=-5‰, and *W*=10 mmol mol^-1^. The values of parameters for simulations were shown in Table S1.

### Overview of g_m_ and g_sc_ variations

Water availability and PFGs significantly impacted *A*_n_, *g*_sc_, and *g*_m_ (Table 3). Compared with the non-stressed condition, drought stress decreased *A*_n_, *g*_sc_, and *g*_m_ for all three PFGs significantly (Fig. 2), but the magnitude of the decrease varied between PFGs. Within a PFG, drought stress reduced *A*_n_ by 36% for the DW, 67% for the EW and 42% for the HB (Fig. 2a); and reduced *g*_sc_ by 37% for the DW, 81% for the EW and 47% for the HB (Fig. 2b), on average. The magnitude of the reduction in *g*_m_ was similar to that of *g*_sc_: 36% in the DW, 80% in the EW and 35% in the HB PFG (Fig. 2c). The HB had the highest values of *A*_n_, *g*_sc_, and *g*_m_ among the three PFGs under both non-stressed and drought-stressed conditions (Fig. 2). That is, the higher *A*_n_, the higher were *g*_sc_ and *g*_m_ within a functional group. This relationship was generally true also for the individual species within a PFG. Accordingly, positive correlations between *A*_n_ and *g*_sc_ or *g*_m_ were found for the different species of a PFG across water stress treatments, although the quantitative relations were significantly different between PFG_S_ (Supplementary Data Fig. S1).

**Table 3.** ANOVA tests for *A*_n_, *g*_sc_, *g*_m_, *g*_sc_/*g*_m_, and iWUE_obs_ using a general linear model. Data are square root transformed to improve the normality before ANOVA tests. Significant effects are shown in bold (*p*<0.05)

| Source | df | $A_n$ | | $g_{sc}$ | | $g_m$ | | $g_{sc}/g_m$ | | $iWUE_{obs}$ | |
| --- | --- | --- | --- | --- | --- | --- | --- | --- | --- | --- | --- |
| | | $F$ | $P$ | $F$ | $P$ | $F$ | $P$ | $F$ | $P$ | $F$ | $P$ |
| Drought stress | 1 | 101.44 | **0.000** | 105.43 | **0.000** | 56.13 | **0.000** | 0.004 | 0.949 | 9.91 | **0.002** |
| PFG | 2 | 50.85 | **0.000** | 31.33 | **0.000** | 30.61 | **0.000** | 1.530 | 0.219 | 0.48 | 0.617 |
| W*PFG | 2 | 3.85 | **0.023** | 6.70 | **0.002** | 4.75 | **0.010** | 0.199 | 0.820 | 0.99 | 0.374 |

**Fig. 2.**
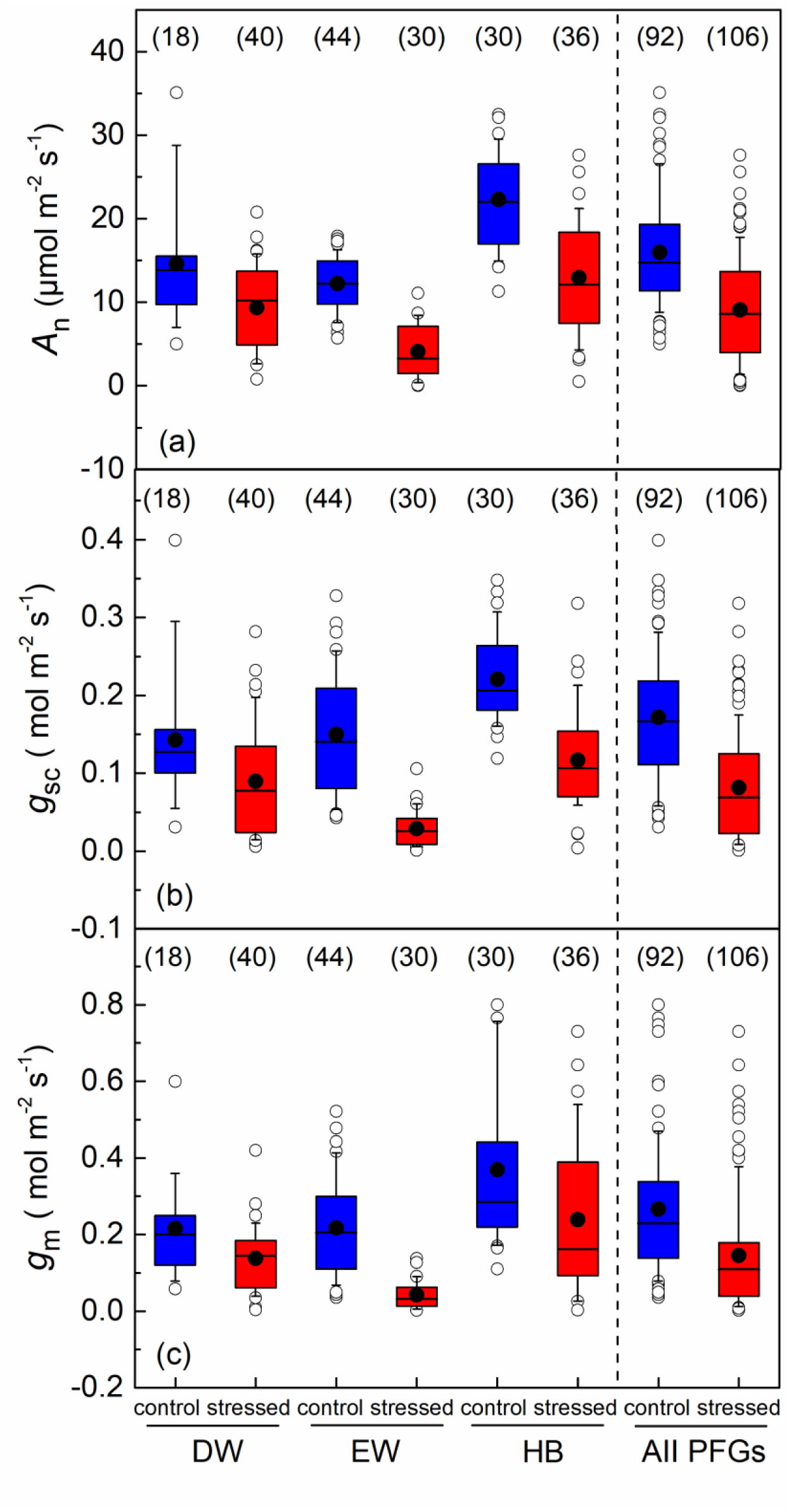
*A*_n_ a), *g*_sc_ b) and *g*_m_ c) across drought-stress levels and plant functional groups (PFGs: DW, deciduous and semi-deciduous woody species; EW evergreen woody species; and HB, herbaceous species). Data collection of different species under non-stressed (blue boxes) or water-stressed conditions (red boxes). Boxplots show median (center line), mean (black dot), inter-quartile range, 10-90% range (whiskers), and outliers (open circles). Numbers in brackets are the number of observations. Data are from the following references: Bongi et al. (1989), Delfine et al. (2001, 2005), Flexas et al. (2002), Galmes et al. (2007), Miyzawa et al. (2008), Barbour and Kaiser (2016), Brilli et al. (2013), Cano et al. (2013, 2014), Centritto et al. (2009), Ferrio et al. (2012), Perez-Martin et al. (2009, 2014), Warren et al. (2008, 2011).

The *g*_sc_/*g*_m_ ratio showed no significant differences between water availability conditions or PFGs (*p*>0.05), while iWUE_obs_ was significantly influenced by drought stress (Table 3). Drought stress improved iWUE_obs_ by 25% compared with non-stress conditions when pooling the data of all plant functional groups (Fig. 3b). The *g*_sc_/*g*_m_ ratio of drought-stressed plants (0.79±0.1 1, 95%CI) was almost identical to that of non-stressed plants (0.79±0. 09, 95%CI) (*p*>0.05) (Fig.3a). The *g*_sc_/*g*_m_ ratio of the EW (0.87±0.13, 95%CI) appeared to be higher, but not significantly so, than that of the DW (0.71±0.11, 95%CI) and the HB (0.77±0.12, 95%CI) PFG. The mean g_sc_/g_m_ ratio was 0.79 (±0.07, 95%CI, n= 198) across drought-stress levels and PFGs. There was a significant correlation between *g*_m_ and *g*_sc_ that was similar for drought-stressed (*r*^2^=0.58, *n*=106, *p*<0.0001, Fig. 4) and non-stressed conditions (*r*^2^=0.20, *n*=92, *p*<0.0001, Fig. 4). When pooling all data of the main dataset to include other long-term treatments, *g*_sc_ and *g*_m_ were also positively correlated (*r*^2^=0.47, *n*=364, *p*<0.0001) with a mean *g*_sc_/*g*_m_ of 0.88±0.06 (95%CI) (Supplementary Data Fig. S2).

**Fig. 3.**
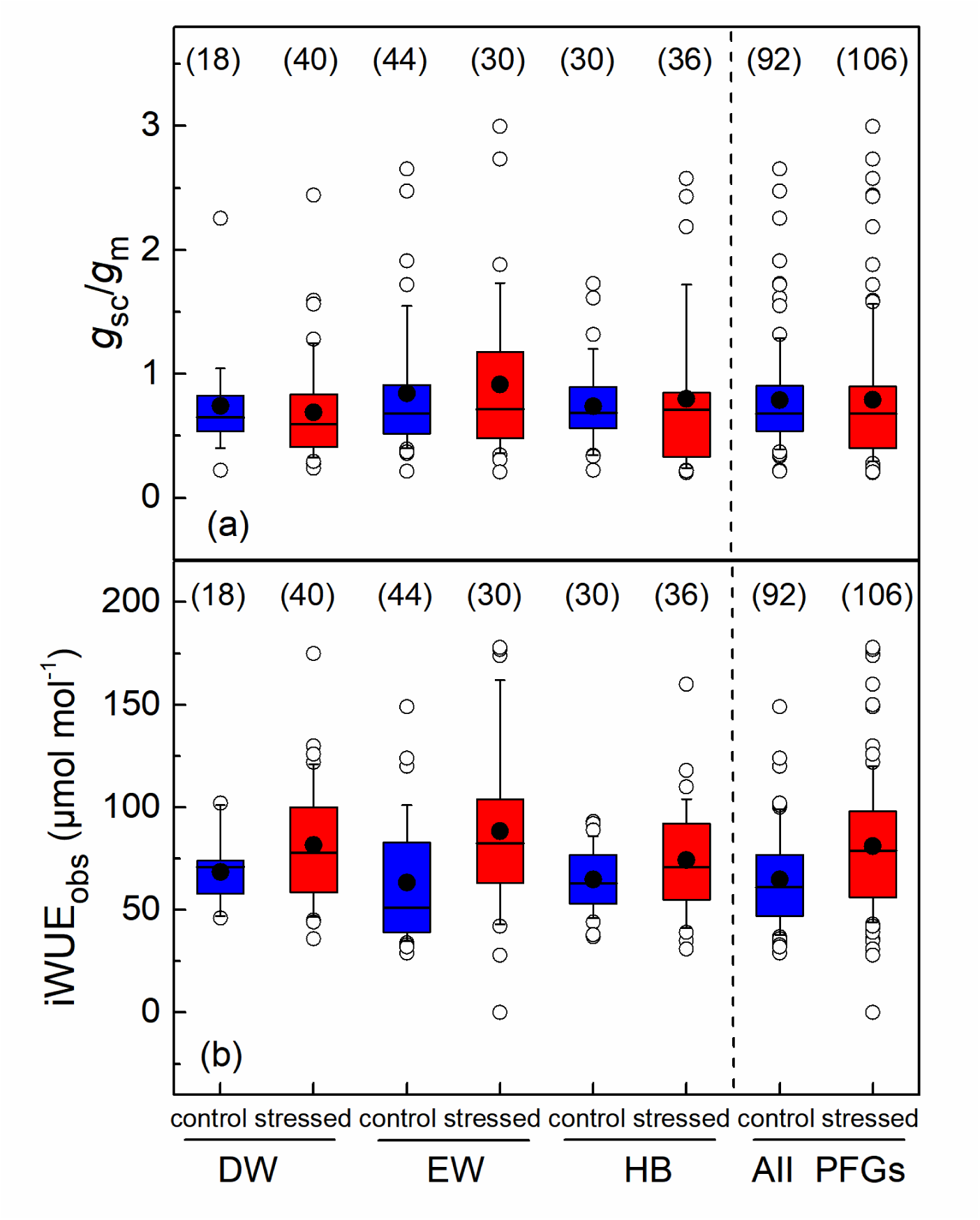
*g*_sc_/*g*_m_ a) and iWUE_obs_ b) across drought treatments and plant functional groups (PFGs: DW, deciduous and semi-deciduous woody species; EW evergreen woody species; and HB, herbaceous species). Data source and the meaning of symbols as in Fig. 2.

**Fig. 4.**
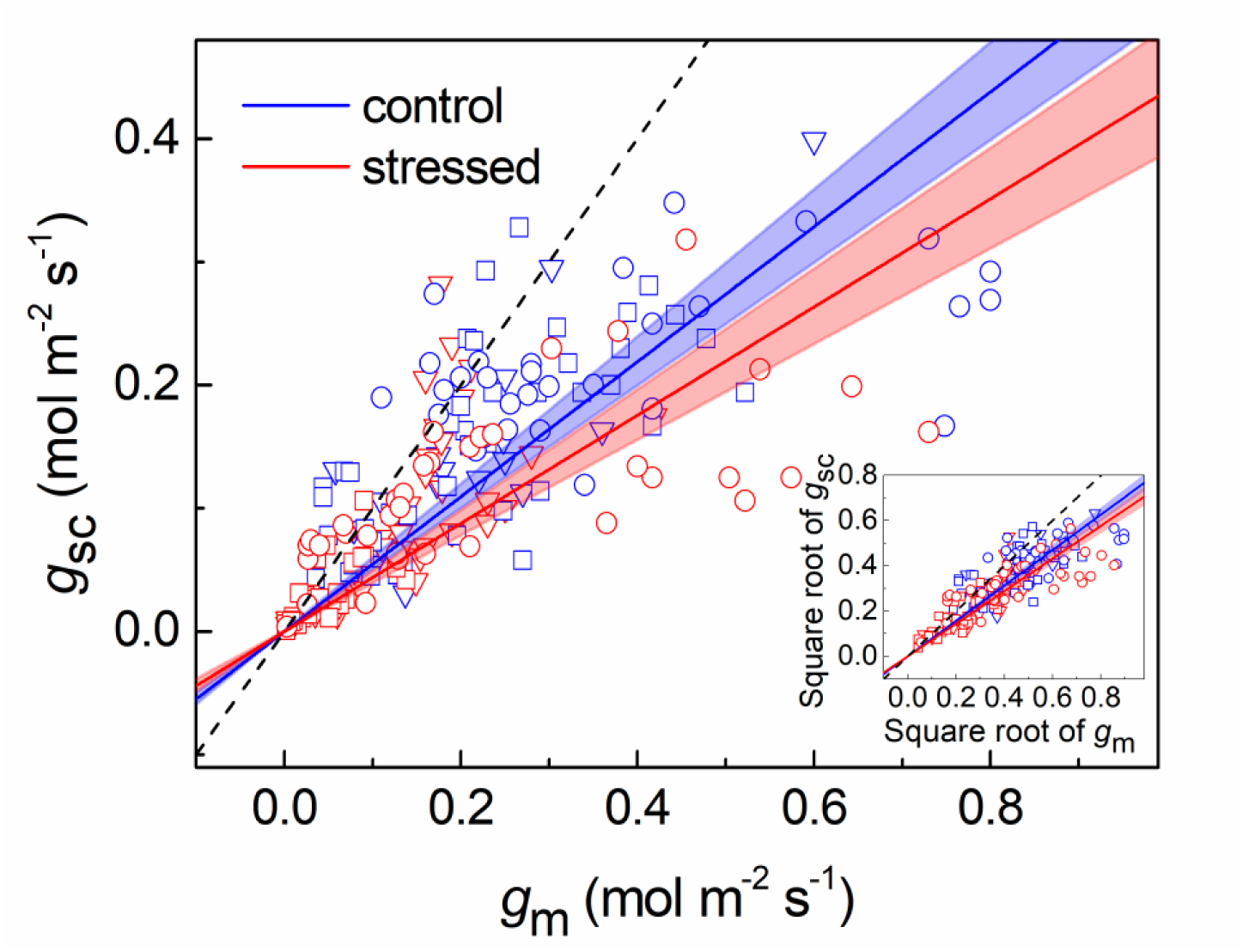
The relation between *g*_m_ and *g*_sc_ of non-stressed plants (blue solid lines) and drought-stressed plants (red solid line) pooling over plant functional groups. Symbols and species: ▽, deciduous semi-deciduous woody species (DW); □, evergreen woody species (EW); ○, herbaceous species (HB). The internal panel shows the relation between square-root transformed *g*_m_ and *g*_sc_. Regressions are linear function through the origin across all genotypes with the slope of 0.78±0.02 (SE) of non-stressed plants (*r*^2^=0.20, *p*<0.0001, *n*=92) and 0.72±0.02 (SE) of drought-stressed plants (*r*^2^=0.58, *p*<0.0001, *n*=106) for the square-root transformed data. The dashed black line is a 1:1 line.

### Predicting iWUE from photosynthetic ^13^C discrimination Δ

A value for *g*_sc_/*g*_m_ of 0.79 was used to compute iWUE_mes_ (Eqn 12) and thus predict iWUE from day-respiration corrected Δ (Δ_P_) in seven species in controlled-environment experiments reported in Table 2. iWUE_mes_ agreed much better with observed values of iWUE_obs_ than iWUE_sim_ as shown by the regression analysis (Fig. 5a). This was further supported by the RMSE which was far lower for iWUE_mes_ than for iWUE_sim_ (Fig. 5). The mean error of iWUE_mes_ (5 μmol mol^-1^) was also significantly lower than that of iWUE_sim_ (53 μmol mol^-1^). The difference in RMSE between the two models was largest (about 62 μmol mol^-1^) in *G. max* and smallest in *H. lanatus* and *T. aestivum*, at only about one half that of *G. max*. The accuracy of predicted iWUE using iWUE_mes_ thus seemed to vary somewhat between species, likely related to interspecific differences in *g*_sc_/*g*_m_.

**Fig. 5.**
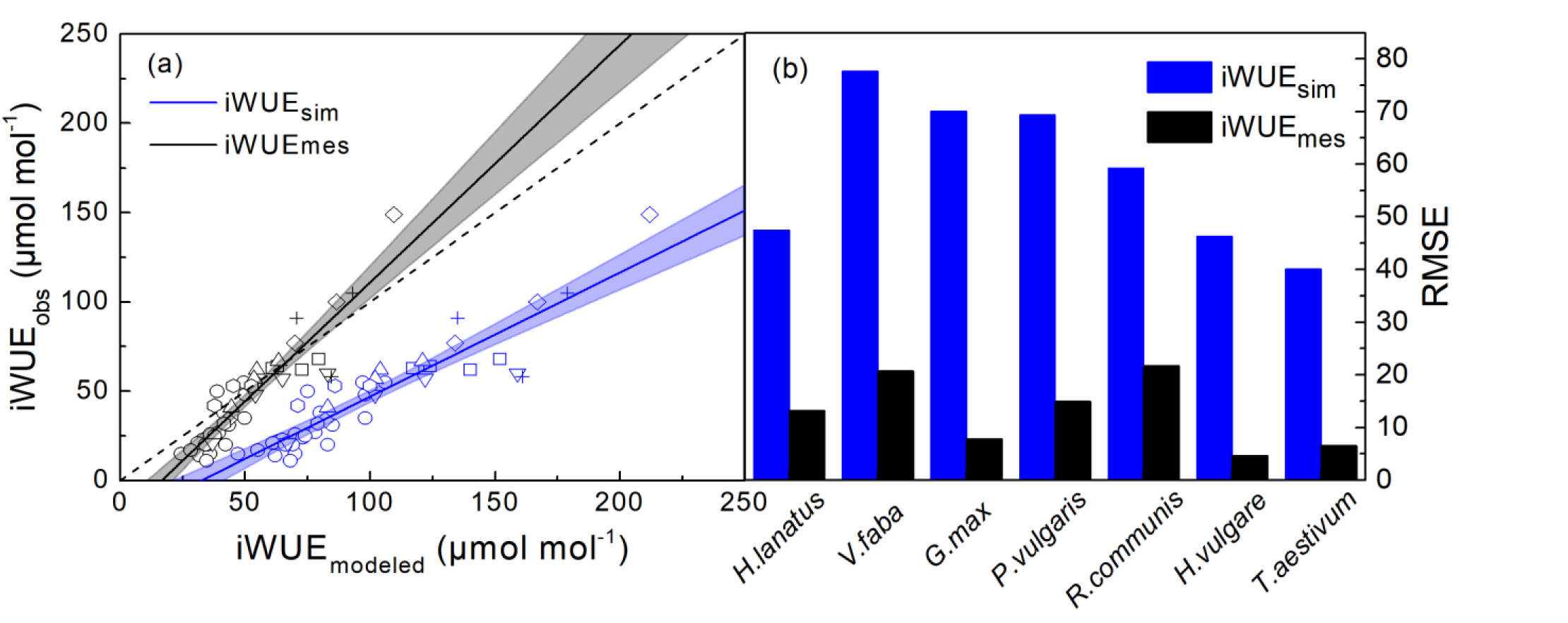
iWUE calculated from photosynthetic Δ using the simple model (iWUE_sim_) and the iWUE_mes_ model compared to the measured iWUE (iWUE_obs_) a) and the comparison of RMSE of the two models b). Symbols and species: □, G. max; ○, *H. lanatus*; △, *H. vulgare*; ▽, *P. vulgaris*; ◊, *R. communis*; 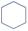, *T. aestivum*; +, *V. faba*. The solid lines are least-squares linear regressions: (a) iWUE_obs_=-22.5(±4.7)+0.7(±0.05)iWUE_sim_, *r*^2^=0.84, *p*<0.0001; (b) iWUE_obs_=-22(±4.7)+1.3 (±0.09)iWUE_mes_, *r*^2^=0.84, *p*<0.0001. Values are mean±SE (*n*=46). The dashed black line is a 1:1 line.

We analyzed the relationship between the measured *g*_m_ or *g*_sc_/*g*_m_ and iWUE_obs_ also in the drought-stress dataset and the two independent datasets, where *g*_m_ was measured using the online Δ method of Gong *et al*. (2015, 2018). This revealed a weak but statistically significant negative correlation between *g*_m_ and iWUE_obs_ across drought-stress levels and species (*p*<0.01, *r*^2^<0.1, Fig. 6a), a correlation that was also confirmed in our experimental datasets (*p*<0.0001, *r*^2^=0.37, Fig. 6b). We found a significant negative correlation between *g*_sc_/*g*_m_ and iWUE_obs_ in all datasets (Fig. 6cd). This relation was more robust than the *g*_m_-iWUE_obs_ relationship as was indicated by the higher *r*^2^, in both the drought-stress dataset (*p*<0.0001, *r*^2^=0.25, Fig. 6c) and the independent experimental datasets (*p*<0.0001, *r*^2^=0.50, Fig. 6d). Beyond that, relations between *g*_sc_/*g*_m_ and iWUE_obs_ were statistically identical (i.e. regression parameters were not significantly different) for both datasets.

**Fig. 6.**
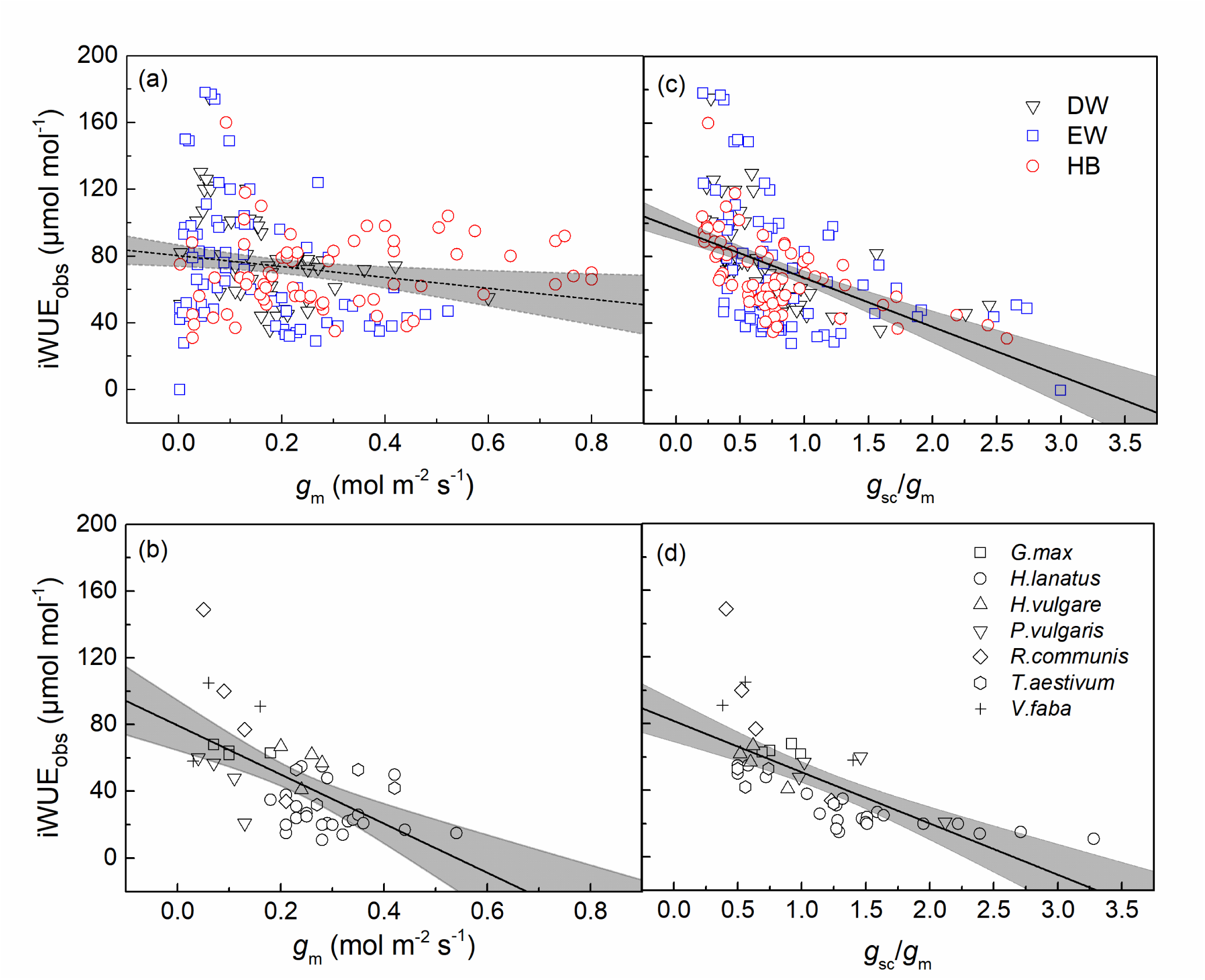
Relation between the measured iWUE (iWUE_obs_) and *g*_m_ of a) synthesized drought-stress dataset and b) experimental dataset of Table 2; The lines are least-squares linear regressions across all species: a) iWUE=80.10(±3.29)-32.56(±12.46)*g*_m_, *r*^2^=0.03, *p*<0.01, *n*=198; b) iWUE=79.52(±7.48)-147.12(±29.03)*g*_m_, *r*^2^=0.37, *p*<0.0001, *n*=46; c) iWUE=96.75(±3.43)-29.39(±3.63)*g*_sc_/*g*_m_, *r*^2^=0.25, *p*<0.0001, *n*=198; d) iWUE=81.67(±6.25)-30.77(±4.69)*g*_sc_/*g*_m_, *r*^2^=0.50, *p*<0.0001, *n*=46; values are mean±SE.

## DISCUSSION

### Including mesophyll conductance in iWUE estimation is critical

The importance of accounting for *g*_m_ in the Δ-iWUE relationship is supported by our model analysis, which illustrates that omitting mesophyll conductance (or the use of iWUE_sim_ which assumes an infinite *g*_m_) causes a significant overestimation of observed iWUE. Of course, the impact of omitting mesophyll conductance is large when mesophyll conductance is small and affecting Δ considerably. At a “common” Δ value of 18‰, the iWUE_sim_ model overestimated iWUE by 65%, clearly showing that iWUE_sim_ did not provide a reasonable estimate of iWUE. The overestimation of iWUE resulting from omission of mesophyll conductance was even greater at a lower Δ, showing that the use of iWUE_sim_ is even more problematic under conditions that lead to decreasing Δ. This means, that caution must be taken when comparing iWUE of plants in conditions that lead to divergent Δ, such as contrasting soil water availability, temperature, or transpiration demand when using the iWUE_sim_ model.

Quantitative prediction of iWUE is essential to anticipate plant responses to climate change factors like CO_2_ and precipitation regime (Adams *et al*., 2019; Soh *et al*., 2019). Recent studies have shown an increased need to quantify the rate at which iWUE has been changing in different geographical regions (Adams *et al*., 2019). Here, using iWUE_mes_ to account for the *g*_sc_/*g*_m_ ratio seems necessary. By contrast, the impact of respiratory fractionation, boundary layer conductance, and ternary effect on iWUE_mes_ values is far less critical. This not only supports assumptions we made to derive iWUE_mes_ (in Eqn 12), but also means that comparing crop varieties for iWUE_mes_ should not be compromised significantly by changes in respiratory metabolism or leaf morphology affecting boundary layer.

An accurate estimation of iWUE is also essential to predict relations between water and carbon fluxes at scales higher than the leaf scale. For instance, combining iWUE estimates from the ^13^C discrimination and transpiration flux inferred from sap flow has been used to quantify photosynthesis of forest canopies (Klein *et al*., 2016). The isotope fractionation can be estimated using the δ^13^C value of either bulk organic matter or carbohydrates (via compound-specific δ^13^C analysis of sugars). The δ^13^C in sugars is believed to be better proxy for iWUE because it reflects recent photosynthetic assimilation and is less influenced by post-photosynthetic fractionation processes (Hobbie & Werner, 2004; Badeck *et al*., 2005). Smith *et al*. (2016) showed that using the δ^13^C of leaf and phloem carbohydrates improved the precision of iWUE estimations, as compared to using the δ^13^C value of bulk soluble carbon. However, both methods overestimated iWUE when iWUE_sim_ was used. We used their data (Smith *et al*., 2016) and calculated iWUE_mes_ using a *g*_sc_/*g*_m_ of 0.79, with the value of Δ obtained from individual carbohydrates (i.e. sucrose, glucose, and fructose). This significantly reduced the error in predicted iWUE (Fig. S3). That is, neglecting the impact of *g*_m_ also overestimates considerably iWUE calculated from the δ^13^C value of individual carbohydrates.

### Parameterization of iWUE_mes_ calculation

As shown by Eqn 11 or 12, calculating iWUE_mes_ from Δ requires the knowledge of the *g*_sc_/*g*_m_ ratio. Our study suggests that a value of *g*_sc_/*g*_m_ of 0.79 can be applied broadly. In fact, that ratio did not seem to be influenced by drought stress or plant functional groups in the global dataset. Using such a value of the *g*_sc_/*g*_m_ ratio can therefore be useful to compare iWUE_mes_ values of multiple species grown under varying soil water conditions, particularly if there is no direct information on mesophyll conductance. The use of a value of 0.79 is further supported by our independent experimental data of seven species (Fig. 5). It is worth noting that even the use of a constant value of 0.79 does not simply change the scale of iWUE values by a proportionality constant, because the term associated with *g*_m_ in iWUE_mes_ is an additive term in the denominator (Eqn 12). The apparent insensitivity of the *g*_sc_/*g*_m_ ratio to drought relates to the similar sensitivity of *g*_sc_ and *g*_m_ to long-term water stress, as both were reduced by about 50% across plant functional groups. In other words, our results do not directly support the hypothesis of a lower sensitivity of *g*_m_ to drought stress compared to *g*_sc,_ to help maintaining leaf carbon gain (Cano *et al*., 2014). Also, *A*_n_ appeared to decline less than *g*_sw_, leading to an increased iWUE under sustained drought-stress, as we previously found in the field (Gong *et al*., 2011). In addition, our compilation of observations on multiple species shows a negative correlation between *g*_m_ and iWUE_obs_ across drought-stress levels, challenging the common assumption that a high *g*_m_ is beneficial to water-use during photosynthesis (Barbour *et al*., 2010; Flexas *et al*., 2013; Barbour & Kaiser, 2016). In other words, our survey suggests that increased iWUE may not necessarily be associated with high *g*_m_.

Although a common *g*_*sc*_*/g*_*m*_ value of 0.79 seems to be an acceptable compromise when information on mesophyll conductances is not available, it is certainly desirable to parameterize iWUE_mes_ with proper experimental values in order to compute precise intrinsic water use efficiency values. In fact, we found a significant, negative relationship between iWUE_obs_ and *g*_sc_/*g*_m_ across a large range of species (Fig. 6). In addition, *g*_sc_/*g*_m_ explained a higher proportion of variation in iWUE_obs_ compared to *g*_m_ alone, as expected from Eqn 12. Therefore, *g*_sc_/*g*_m_ contributed to explaining better the observed interspecific difference in iWUE_obs_. That is also probably the case between varieties within the same species, and thus the *g*_sc_*/g*_m_ ratio should be considered in breeding programs targeting water-use efficiency.

We nevertheless recognize that variations in *g*_sc_/*g*_m_ amongst functionally similar species or between studies is the main challenge in the use of iWUE_mes_. Current studies show that responses of *g*_m_ to environmental factors are highly complex and such factors include CO_2_ (Flexas *et al*., 2007), temperature (von Caemmerer & Evans, 2013; Shrestha *et al*., 2019) and VPD (Loucos *et al*., 2017, Stangl *et al*., 2019). Up to now, variations in *g*_m_ cannot be easily related to anatomical features and it is believed that biochemical processes (e.g. facilitated diffusion by carbonic anhydrase or aquaporins) are involved (Flexas *et al*., 2008) although controversial results have been obtained (Perez-Martin *et al*., 2014; Kromdijk *et al*., 2020). It is also worth noting that observed variations in *g*_m_ could be related to the methods used for measurements. Some authors questioned the robustness of current observed values of *g*_m_ (Tholen *et al*., 2012; Sun *et al*., 2014b), because current methods extract *g*_m_ from a residual photosynthetic component after other processes have been accounted for, meaning that all errors (artifacts) accumulate in the estimate of *g*_m_ (Pons *et al*., 2009). For example, Gong *et al*. (2015) demonstrated that *g*_m_ estimates can be strongly influenced by diffusive leaks and isotopic disequilibria between respiration and photosynthesis when *g*_*m*_ is computed from non-corrected online Δ values. The contribution of the apparent isotopic fractionation by day respiration could easily be misquantified if carbon reserves are respired instead of current photosynthates, thus causing errors in *g*_m_ estimates (Gong *et al*., 2015; Tcherkez *et al*., 2017; Barbour *et al*., 2017; Busch *et al*., 2020). The use of iWUE_mes_ effectively provides much more accurate iWUE values than iWUE_sim_, but might be associated with an error (e.g. for species with a different *g*_sc_/*g*_m_ ratio) if the ‘standard’ 0.79 *g*_sc_*/g*_m_ value is used. Certainly, in mechanistic studies of the controls of iWUE in a given species it is warranted to examine environmental effects on relationships between *g*_sc_ and *g*_m_ with experimental detail.

## Perspectives

Our analysis shows that the expression of iWUE_mes_ that accounts for mesophyll conductance (Eqn 12) can be used to provide better estimates of intrinsic water use efficiency. Together with an observation-based *g*_sc_/*g*_m_ ratio, it can be used to reanalyze, adjust or correct published iWUE_sim_ data for which measurements of *g*_m_ are unavailable. This has far reaching importance, as *g*_m_ is a key parameter for most of the models of leaf photosynthesis or carbon cycling, but is often poorly represented in them (Tholen *et al*., 2012; Sun *et al*., 2014b). Assuming an infinite *g*_m_ leads to an overestimation of global gross primary production by about 16% by earth system models (Sun *et al*., 2014a). Scaling of *g*_m_ to *g*_sc_ is potentially a useful way to reduce the error related to *g*_m_ in such models. Our study also examined potential differences in *g*_sc_/*g*_m_ ratios across long-term drought-stress treatments and plant functional groups and showed that *g*_sc_/*g*_m_ has rather similar values such that an average value of 0.79 in the iWUE_mes_ model works well to provide more realistic iWUE values. However, in an effort to compare and select varieties based on water use efficiency, genotypic variation in *g*_sc_*/g*_m_ should not be overlooked. That is, the effect of mesophyll conductance (relative to that of stomatal conductance) must be accounted for to provide reliable estimates of iWUE based on carbon isotopes. This is particularly true in crop species where mesophyll conductance contributes significantly to photosynthetic limitations such as *V. faba, G. max* and *P. vulgaris* in this study.

## ACKNOWLEDGMENTS

This work was supported by the National Natural Science Foundation of China (NSFC 31870377), the Guangdong Natural Science Foundation (2018A030313450) and the Deutsche Forschungsgemeinschaft (SCHN 557/7-1 and SCHN 557/9-1).

## AUTHOR CONTRIBUTIONS

X.Y.G. designed and planned the research; G.T. proposed the iWUE_com_ model and the framework of sensitivity analyses; W.T.M. performed the literature survey and data analyses; W.T.M., X.M.W and X.Y.G. wrote the first draft, and all authors discussed the results and implications and contributed to the revision.

## Supporting Information

**Fig. S1.**
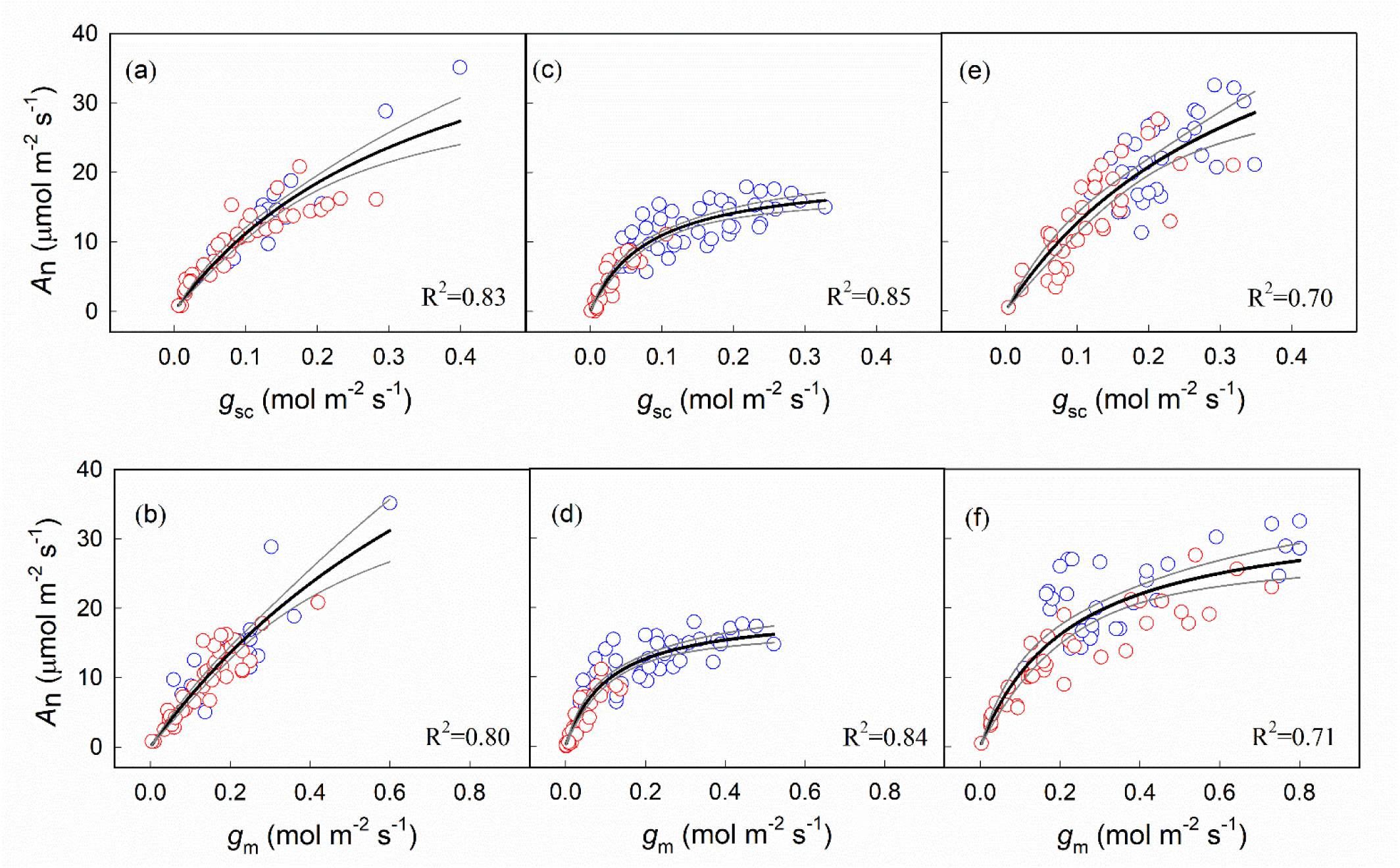
Relationships between *A*_n_ and *g*_sc_, or *g*_m_ across two water availability levels (control, blue dots and drought stress, red dots) for the DW (a and b), EW (c and d), and HB (e and f). Data were fitted use a function of y= *a*x/(*b*+x), *n*=58 for DW, *n*=74 for EW, *n*=66 for HB, and *p*<0.0001 for all regressions.

**Fig. S2.**
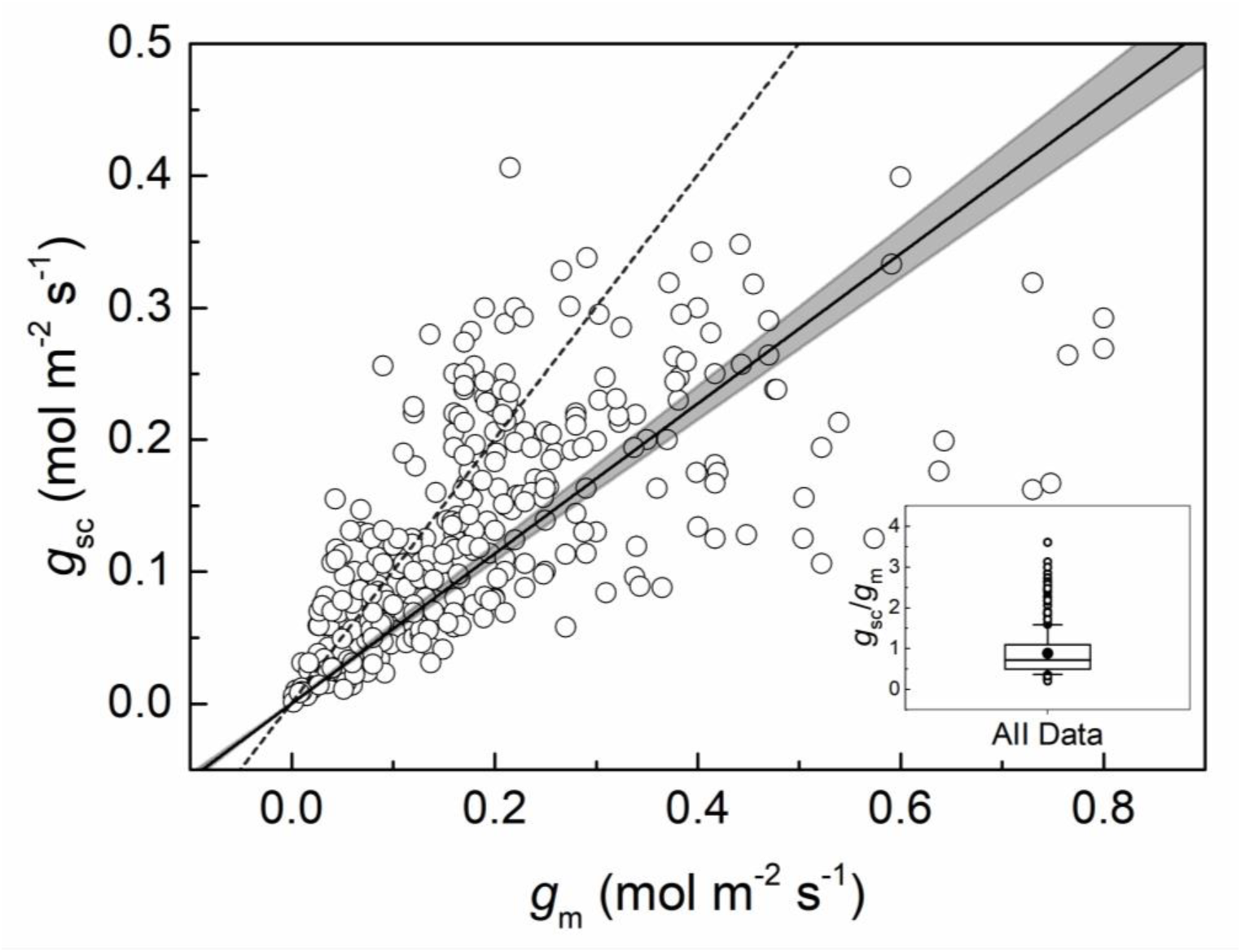
The relation between *g*_m_ and *g*_sc_ across treatments and species in the main dataset compiled from the published papers. The internal panel shows a box plot of *g*_sc_/*g*_m_ calculated from the same data with a mean *g*_sc_/*g*_m_=0.88(±0.06 95%CI). The dashed line is a 1:1 line, and the solid line is the regression line: *r*^2^=0.47, *p*<0.0001, *n*=364.

**Fig. S3.**
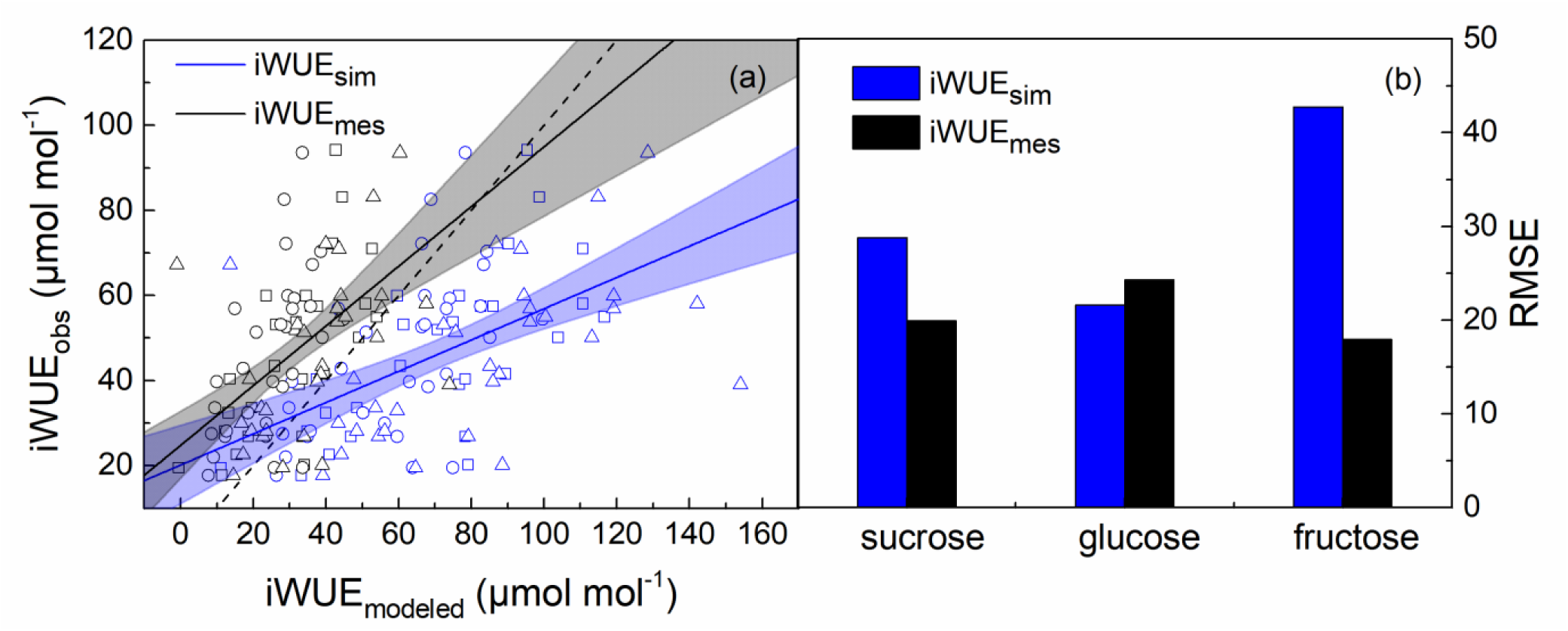
Comparison of the measured iWUE_obs_ and the modeled iWUE using the simple (iWUE_sim_, blue symbols and lines) and the iWUE_mes_ model (black symbols and lines) calculated from the data of Smith *et al*. (2016) using Eqn 13. Symbols and leaf metabolites: □, sucrose; ○, glucose; △, fructose. The dashed black line is a 1:1 line. The solid lines are least-squares linear regressions: iWUE_obs_=20.17(±4.63)+0.37(±0.06)iWUE_sim_, *r*^2^=0.30, *p*<0.0001; iWUE_obs_=24.57(±3.97)+0.69(±0.11)iWUE_mes_, *r*^2^=0.30, *p*<0.0001. Values are mean±SE (*n*=90).

**Fig. S4.**
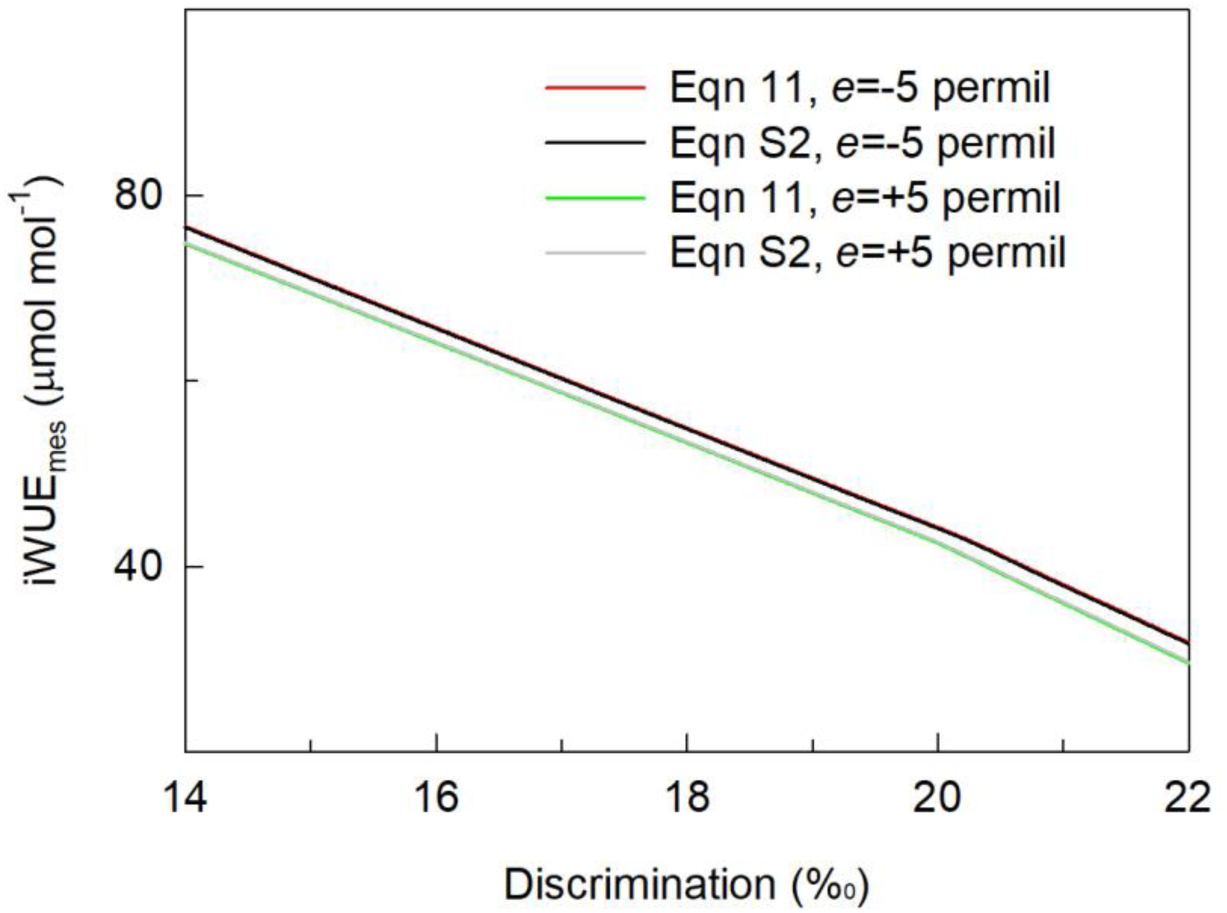
Comparison of the iWUE_mes_ prediction estimated from Eqn 11 and S2 as affected by the influence of mitochondrial respiratory fractionation (*e*)

**Table S1.**
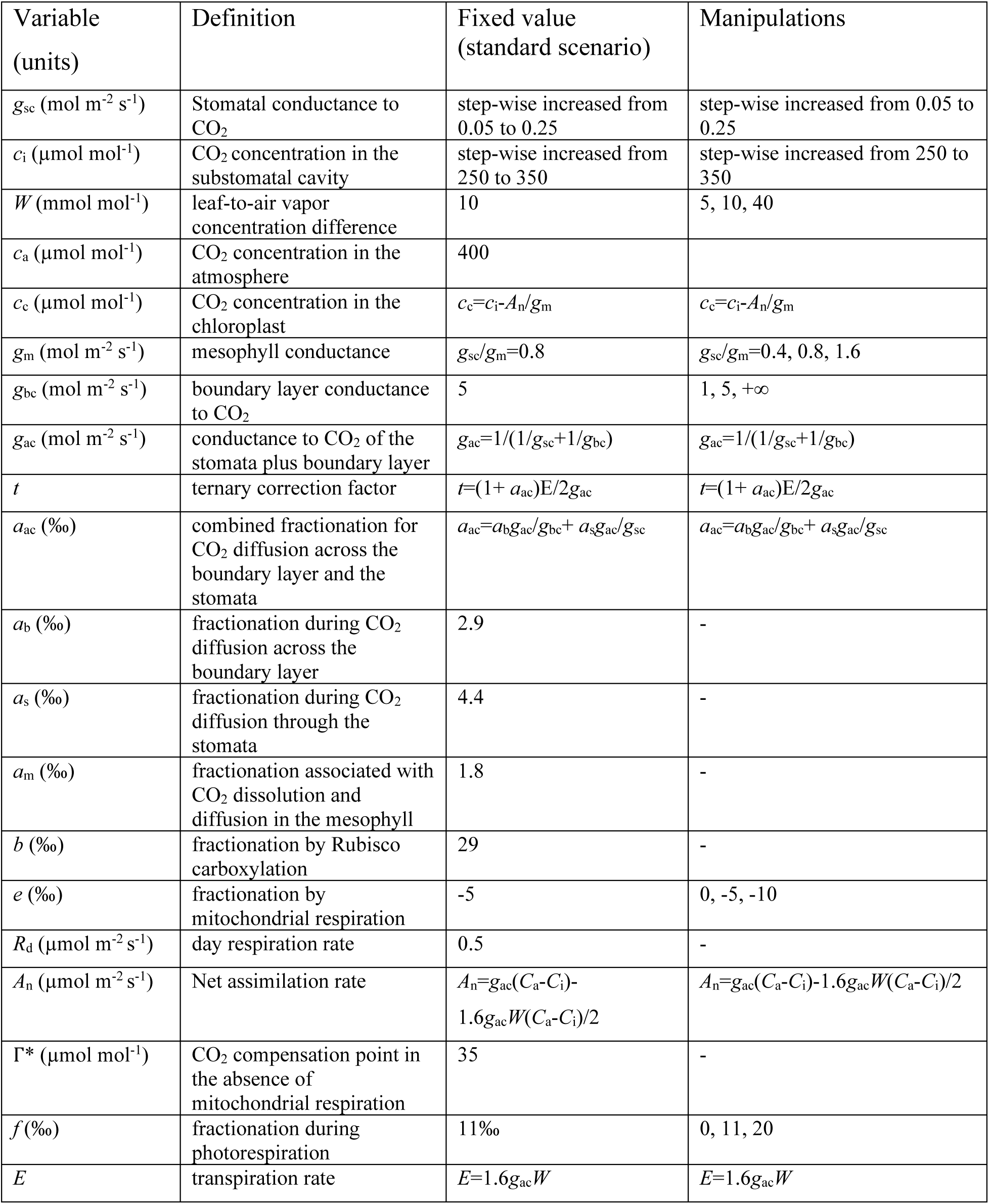
Variables used for sensitivity tests. *g*_sc_ and *c*_i_ were stepwise increased to create a gradient of photosynthetic discrimination, and iWUE models were used to calculate iWUE_sim_, iWUE_mes_, and iWUE_com_.

**Note S1** Derivation of Eqn 8 from the comprehensive photosynthetic discrimination model. Using the notations *e’* = *eα*_*b*_*/α*_*e*_, *f’* = *fα*_*b*_*/α*_*f*_, and substituting *c*_c_ by *c*_i_ – *A*_n_*/g*_m_, Eqn 3 gives:

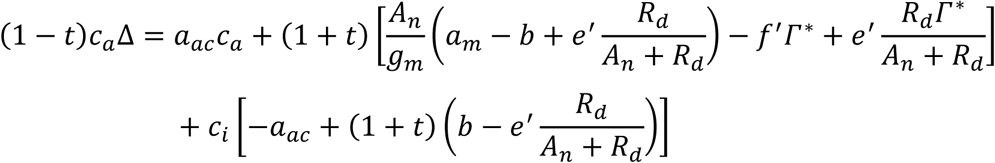

Rewriting *A*_*n*_*/g*_*m*_ as (*A*_*n*_/*kg*_*ac*_)*kg*_*ac*_*/g*_*m*_ = iWUE^*^_com_ *kg*_*ac*_*/g*_*m*_ and using Eqn 7 leads to:

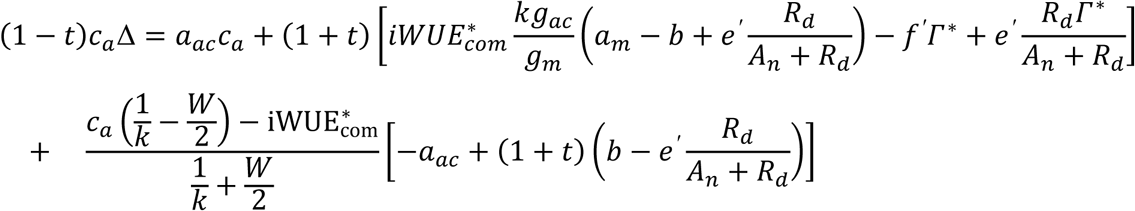

Rearranging (factorization by iWUE^*^ _com_) gives:

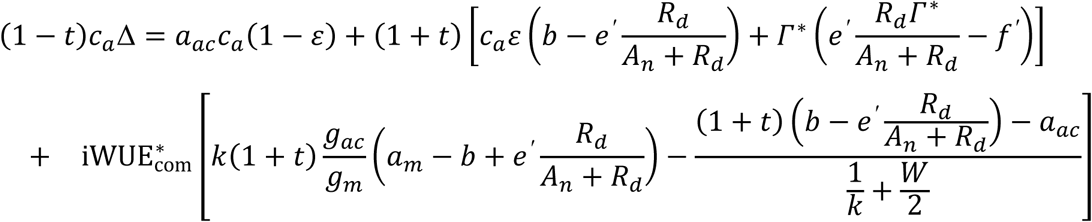

where *ε* is defined by Eqn 9. Rearranging to express iWUE^*^_com_ gives Eqn 8).

**Note S2 The modified carbon isotope discrimination model for partitioning day respiration and its isotopic contribution to photosynthetic discrimination**

The comprehensive model of Eqn 3 can be modified as:

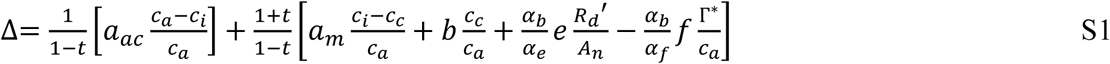

by assuming that day respiration is separated from the pool of primary assimilates during a short exposure to an atmosphere with a different δ^13^C value in CO_2_. Furthermore, *R*_d_*’* is defined as the respired CO_2_ that is released into the ambient air, thus is a net flux whereas *R*_d_ in the original Farquhar model was defined as a gross respiratory flux (net and gross referring here to refixation) (Tcherkez et al. 2017). That is, *R*_*d*_*’* is considered as the flux of CO_2_ liberated by the leaf from respiratory substrates with a slow turn-over and thus not influenced by the isotope composition of photosynthetic assimilates in the short term. Using Eqn S1, *R*_d_’ can be estimated from measurements of online isotope discrimination in a gas exchange system where inlet CO_2_ can be switched between two sources of CO_2_ with contrasted δ^13^C values. More details are provided in Gong et al. (2015; 2018).

We can then derive equations for iWUE similarly as in Note S1. Using the notations *e’* = *eα*_*b*_*/α*_*e*_, *f’* = *fα*_*b*_*/α*_*f*_, and substituting *c*_c_ by *c*_i_ – *A*_n_*/g*_m_, Eqn S1 gives:

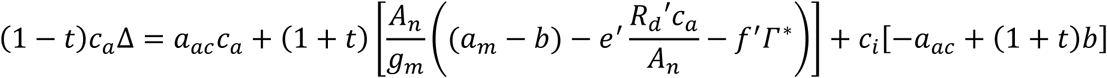

Rewriting *A*_*n*_*/g*_*m*_ as (*A*_*n*_/*kg*_*ac*_)×*kg*_*ac*_*/g*_*m*_ = iWUE^’^_com_ *kg*_*ac*_*/g*_*m*_ and using Eqn 7 leads to (the symbol prime for iWUE refers to the fact we now use a “disconnected” respiration term):

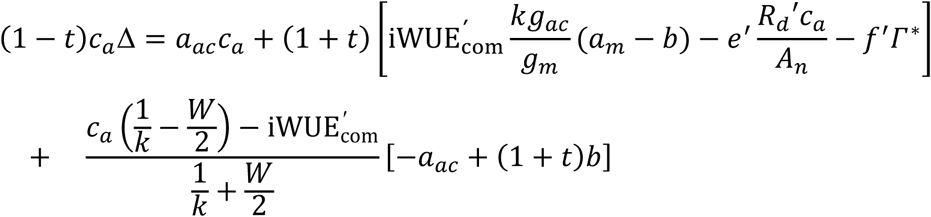

Rearranging (factorization by iWUE^’^_com_) gives:

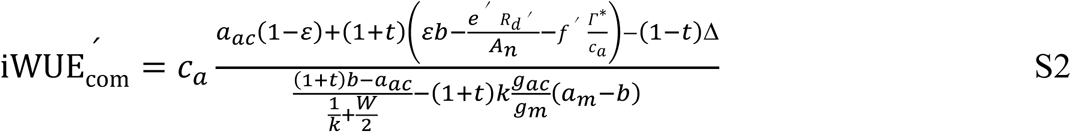

Again, ignoring ternary corrections and boundary layer resistance (*t*=0, *ε*=1, *c*_a_=*c*_s_, *g*_as_=*g*_sc_, *a*_ac_= *a*_s_), Eqn S2 simplifies to:

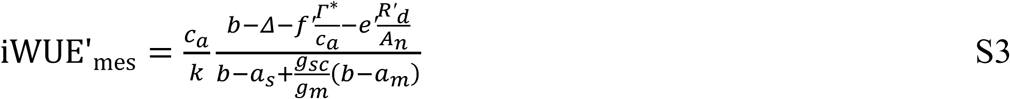

If we further neglect the term associated with day respiration, Eqn S3 simplifies to Eqn 12. Using a series of theoretical data of the standard scenario (Table S1, *g*_sc_/*g*_m_=0.8), we calculated iWUE’_mes_ (Eqn S3) and compared with iWUE_mes_ (Eqn 11), showing that the estimates provided by the two equations (which account for the respiratory fractionation term differently) were very similar (the difference being less than 1μmol mol^-1^, see below). Taken as a whole, within a range of intrinsic respiratory fractionation *e* (far from very high, artificial isotopic disequilibrium), day respiration has little effect on iWUE_mes_ estimates computed from Eqn 11 and S2 (Fig. S4), and of course have no effect at all on iWUE_mes_ estimates generated by Eqn 12 which neglects the respiratory isotope contribution.

